# Recurrently deregulated lncRNAs associated with HCC tumorigenesis and metastasis revealed by genomic, epigenomic, and transcriptomic profiling in paired primary tumor and PVTT samples

**DOI:** 10.1101/050377

**Authors:** Yang Yang, Lei Chen, Jin Gu, Hanshuo Zhang, Jiapei Yuan, Qiuyu Lian, Guishuai Lv, Siqi Wang, Yang Wu, Yu-Cheng T. Yang, Dongfang Wang, Yang Liu, Jing Tang, Guijuan Luo, Yang Li, Long Hu, Xinbao Sun, Dong Wang, Mingzhou Guo, Qiaoran Xi, Jianzhong Xi, Hongyang Wang, Michael Q. Zhang, Zhi John Lu

## Abstract

Hepatocellular carcinoma (HCC) are highly potent to invade the portal venous system and subsequently develop into the portal vein tumor thrombosis (PVTT). PVTT could induce intrahepatic metastasis, which is closely associated with poor prognosis. A comprehensive systematic characterization of long noncoding RNAs (lncRNAs) associated with HCC metastasis has not been reported. Here, we first assayed 60 clinical samples (matched primary tumor, adjacent normal tissue, and PVTT) from 20 HCC patients using total RNA sequencing. We identified and characterized 8,603 novel lncRNAs from 9.6 billion sequenced reads, indicating specific expression of these lncRNAs in our samples. On the other hand, the expression patterns of 3,212 known and novel recurrently deregulated lncRNAs (in >=20% of our patients) were well correlated with clinical data in a TCGA cohort and published liver cancer data. Some lncRNAs (e.g., RP11-166D19.1/MIR100HG) were shown to be useful as putative biomarkers for prognosis and metastasis. Moreover, matched array data from 60 samples showed that copy number variations (CNVs) and alterations in DNA methylation contributed to the observed recurrent deregulation of 716 lncRNAs. Subsequently, using a coding-noncoding co-expression network, we found that many recurrently deregulated lncRNAs were enriched in clusters of genes related to cell adhesion, immune response, and metabolic processes. Candidate lncRNAs related to metastasis, such as HAND2-AS1, were further validated using RNAi-based loss-of-function assays. The results of our integrative analysis provide a valuable resource regarding functional lncRNAs and novel biomarkers associated with HCC tumorigenesis and metastasis.

## Introduction

Hepatocellular carcinoma (HCC) is one of the most common and aggressive human malignancies worldwide (Jemal et al. 2011). The dismal clinical outcome of HCC is largely due to the high incidence of intrahepatic and extrahepatic metastasis in HCC patients (Aldrighetti et al. 2009). HCC cells are highly likely to develop into portal vein tumor thrombosis (PVTT), which is the main route for intra-hepatic metastasis of HCC (Mitsunobu et al. 1996). Therefore, PVTT is closely associated with poor prognosis for HCC patients (Uka et al. 2007). PVTT was found in more than 40% of newly diagnosed HCC cases (Shi et al. 2011), of which the median survival time was less than 12 months with surgical treatment (Minagawa and Makuuchi 2006).

Several long noncoding RNAs (lncRNAs), including MALAT1 (Lai et al. 2012; Gutschner et al. 2013), H19 (Li et al. 2014), HOTAIR (Gupta et al. 2010), and HULC (Panzitt et al. 2007), are directly involved in tumorigenesis and metastasis of various types of cancer. Recent studies have also revealed the pro-metastasis mechanisms through which some lncRNAs contribute to the activation of epithelial-to-mesenchymal transition (EMT) networks, including activation of the WNT (Lau et al. 2014) and TGF-β signaling pathways (Yuan et al. 2014). Although several studies have assessed the contributions of individual lncRNAs to the development of HCC, the functions and mechanisms of only a few individual lncRNAs in HCC tumorigenesis and metastasis are understood in detail (Huang et al. 2007; Braconi et al. 2011; Quagliata et al. 2014; Wang et al. 2014).

Moreover, efforts at systematic identification and characterization of novel lncRNAs involved in HCC, especially those involved in HCC metastasis, remain at an early stage. A recent study based on The Cancer Genome Atlas (TCGA, http://cancergenome.nih.gov/) revealed the existence of more than 50,000 lncRNAs (designated MiTranscriptome lncRNAs) in the human transcriptome (generated from various tumors, normal tissues, and cell lines) (Iyer et al. 2015), of which more than 80% were not reported in previous studies or found in databases (i.e., GENCODE (Harrow et al. 2012) and Refseq (Pruitt et al. 2014)). This study demonstrates the genomic diversity and expression specificity of lncRNAs, while suggesting that more novel lncRNAs will be discovered as additional tumor and cell types (e.g., metastatic samples) are sequenced.

Remarkably, studies suggest that approximately 88% of single-nucleotide polymorphisms (SNPs) in the human genome are within noncoding regions (Maurano et al. 2012), suggesting that many noncoding RNAs and DNA regulatory elements (e.g., promoters and enhancers) have functional roles. Indeed, some lncRNAs play important roles in diverse cellular process, such as cell differentiation (Fatica and Bozzoni 2014), cell death, and tumorigenesis (Du et al. 2013). In addition, lncRNAs can be used as biomarkers for cancer diagnosis, prognosis, and classification because they have cell-type specificity better than that of most protein coding genes and relatively stable local secondary structures, facilitating their detection in body fluids (Cabili et al. 2011; Jun et al. 2015; Yan et al. 2015). For instance, a lncRNA, PVT1, has been used as diagnostic and prognostic biomarker for HCC (Wang et al. 2014).

Here, sixty matched samples (primary tumor, PVTT, and adjacent normal tissue) from 20 Chinese HCC patients were subjected to total RNA-seq (rRNA depleted), followed by integrative analysis at the genomic, transcriptomic, and epigenomic levels, with the goal of identifying and characterizing deregulated lncRNAs in HCC patients. Approximately 76% of the lncRNAs identified in the samples were not annotated by the MiTranscriptome (Iyer et al. 2015) or GENCODE transcriptome (Harrow et al. 2012) databases. Next, approximately 3,000 lncRNAs that were recurrently (in at least 20% of patients) deregulated in primary tumors and/or PVTTs were identified. Their expression levels were correlated with TCGA clinical data and other published liver cancer data (Hong et al. 2015). We also showed that one of the recurrently deregulated lncRNAs was suitable as a prognosis and metastasis biomarker in HCC patients. Furthermore, the aberrant expression patterns of 399 and 342 recurrently deregulated lncRNAs were shown to be due to copy number variations and DNA methylation alterations, respectively. Finally, a coding-noncoding co-expression network was used to predict candidate lncRNAs related to metastasis, after which the predictions were validated experimentally.

## Results

### Identification of novel lncRNAs in 60 matched HCC samples

To systematically identify lncRNAs related to HCC tumorigenesis and metastasis, approximately 9.6 billion reads for 60 samples from 20 HCC patients were sequenced using total RNA-seq (rRNA depleted) (Supplementary File 1). Three matched samples were collected from each patient: primary tumor, adjacent normal tissue, and PVTT.

Novel lncRNAs (Supplementary File 2) were identified using a bioinformatics pipeline (Methods, Figure 1A and Supplementary Table 1). It was based on our *RNAfeature* model (Hu et al. 2015), which was used to predict novel noncoding RNAs in worm (Gerstein et al. 2010), fly, human (Gerstein et al. 2014) and *Arabidopsis* (Di et al. 2014) by integrating expression, RNA secondary structure, conservation and epigenetic signals. Using 13,870 lncRNAs (including 23,898 transcripts) annotated in the GENCODE database (Harrow et al. 2012) (release 19) as a reference, we identified 8,603 novel lncRNAs (including 10,196 transcripts) from the HCC samples, only a small portion of which were reported in other studies. For example, 76% of the newly assembled lncRNAs (we named the identified novel lncRNAs as newly assembled lncRNAs) were not found in the MiTranscriptome database (Iyer et al. 2015), which was mainly derived from TCGA data (Figure 1B). The newly assembled lncRNAs were also compared with two other representative annotation databases: a high quality set, RefSeq (Pruitt et al. 2014), and a comprehensive set, NONCODE (over 50,000 lncRNA transcripts included) (Liu et al. 2005). Only 2% and 10% of the newly assembled lncRNAs were found in the RefSeq (Release 72) and NONCODE (V4) databases, respectively (Supplementary Figure 1). These data indicate the expression specificity of the lncRNAs identified in our samples. In addition, the total RNA-seq libraries used in this study allowed detection of a large number of novel lncRNA transcripts in comparison with other studies (e.g., current TCGA data used polyA-enriched RNA-seq libraries). We showed that the number of lncRNAs increased when the number of sequenced samples was increased (Supplementary Figure 2), while the detection ability was saturated at ~10 and ~20 out of 60 samples for protein-coding genes and GENCODE lncRNAs, respectively.

**Figure 1.**
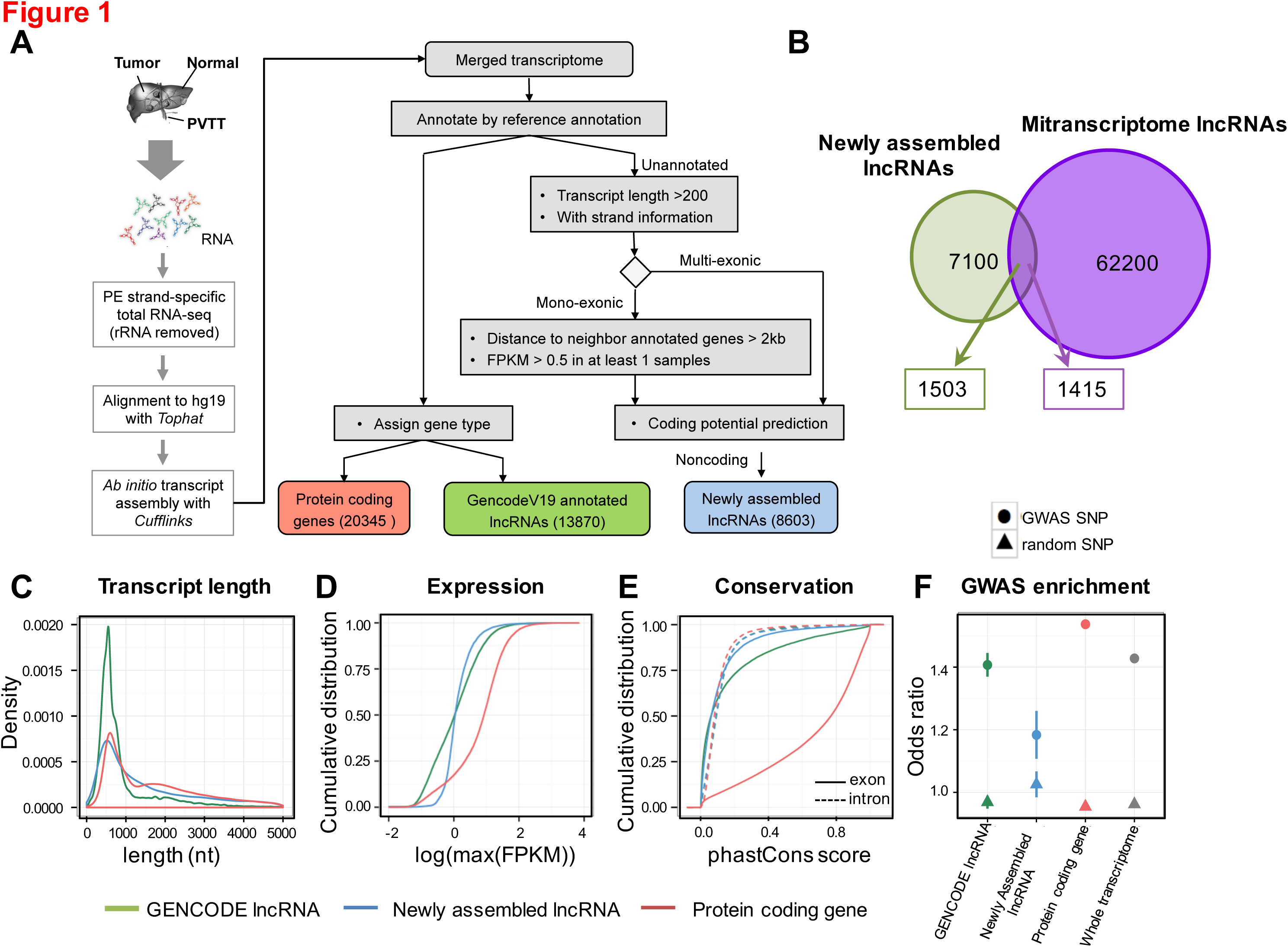
Identify novel lncRNAs in 60 HCC samples. (**A**) Overview of the comprehensive experimental and computational scheme for the systematic identification of lncRNAs in samples from HCC patients.(**B**) Venn diagram showing the fraction of newly assembled lncRNAs that were previously annotated and their overlap with annotations from the MiTranscriptome lncRNA set. Characterization of GENCODE lncRNAs, newly assembled lncRNAs, and protein-coding genes: (**C**) transcript length distribution; (**D**) cumulative distribution curve of maximum gene expression level (RPKM); (**E**) conservation of exons and introns; (**F**) enrichment of GWAS SNPs (circle) over randomly selected SNPs (triangle).

### Characterization of the newly assembled lncRNAs in HCC samples

We characterized genomic location, expression abundance, transcript length, conservation, and SNP enrichment for the newly assembled lncRNAs (Figure 1C-F, Supplementary File 3 and 4). We first focused on the genomic locations of newly assembled lncRNAs (Supplementary Figure 3A). The majority (74%) of the lncRNAs were located in intergenic regions; 16% were antisense to protein-coding genes and 3% were located in introns of protein-coding genes. Moreover, 1.39% and 24.94% of the lncRNAs overlapped with pseudogenes and transposable elements, respectively, whereas 2.24% and 5.06% of the lncRNAs contained local domains conserved with canonical ncRNAs (e.g., rRNA, tRNA, and snRNA) at the sequence and structure levels, respectively. Similarly, the GENCODE lncRNAs also overlapped with or included these elements (Supplementary Figure 3B,C).

Next, we characterized the basic features of the newly assembled lncRNAs and compared them with protein-coding genes and GENCODE lncRNAs. Because they had fewer putative exons, we found that the newly assembled lncRNA transcripts were shorter than protein-coding genes, but longer than GENCODE lncRNAs (Figure 1C, Supplementary Figure 4). This result indicates that the high sequencing depth of our analysis (Supplementary File 1) enables us to assemble long transcripts close to their full length, even though they were expressed at low levels (Figure 1D). Notably, the newly assembled lncRNAs were less evolutionarily conserved in comparison with protein-coding genes and GENCODE lncRNAs, while exonic regions were more conserved in comparison with intronic regions (Figure 1E).

A previous study reported that almost 90% of SNPs are located in non-coding regions (Maurano et al. 2012). To investigate the relationship between lncRNAs and diseases, we capitalized on the GWAS SNP Catalog from NHGRI (Welter et al. 2014). We intersected the lncRNAs with GWAS SNPs from the NHGRI catalog and randomly selected SNPs from the dbSNP (Sherry et al. 2001). Interestingly, we found that GWAS SNPs were significantly enriched in the newly assembled lncRNAs in comparison with a random set (Figure 1F). These data suggested that our newly assembled lncRNAs were likely to be functionally associated with human diseases.

To further validate the activity of the newly assembled lncRNAs, we used published ChIP-seq data for the HepG2 cell line (The ENCODE Project Consortium 2012) to depict activity markers around transcription start sites (TSSs). Different epigenetic signatures (H3K4me3, H3K27ac, Pol II binding, DNase I hypersensitivity) indicate active transcription of the newly assembled lncRNAs in liver cancer cell lines (Supplementary Figure 5). Peaks of these markers were found at the TSSs of lncRNAs, suggesting that the promoters of these transcripts are actively regulated in HepG2 cells.

### Recurrently deregulated lncRNAs in primary tumor and PVTT samples

To identify potential tumorigenesis-and metastasis-associated lncRNAs in HCC samples, we first focused on differentially expressed lncRNAs (including both GENCODE-annotated lncRNAs and newly assembled lncRNAs) for each individual patient between primary tumor tissue and adjacent normal tissue, as well as between PVTT samples and primary tumor cells. We found that the numbers of differentially expressed lncRNAs varied across different patients (Figure 2 and Supplementary Figure 6), especially when comparing PVTT samples with primary tumor samples (Figure 2B and D). In total, we found that 3,203 lncRNAs were recurrently differentially expressed in at least 4 of 20 patients, when comparing primary tumor samples with adjacent normal tissue samples (Figure 2A), whereas only 107 were repeatedly detected when PVTT samples were compared with matched primary tumor samples (Figure 2B). The two sets of recurrently deregulated lncRNAs were designated as tumorigenesis-associated (primary tumor versus adjacent normal tissue) (Figure 2A) and metastasis-associated (PVTT versus primary tumor) lncRNAs (Figure 2B), which are potentially involved in tumorigenesis or metastasis, respectively. The recurrently deregulated lncRNAs are provided in the supplementary material (Supplementary File 6). Notably, of the 107 metastasis-associated lncRNAs, 98 were also recurrently deregulated in primary tumor samples versus paired normal tissues (see details in Discussion).

**Figure 2.**
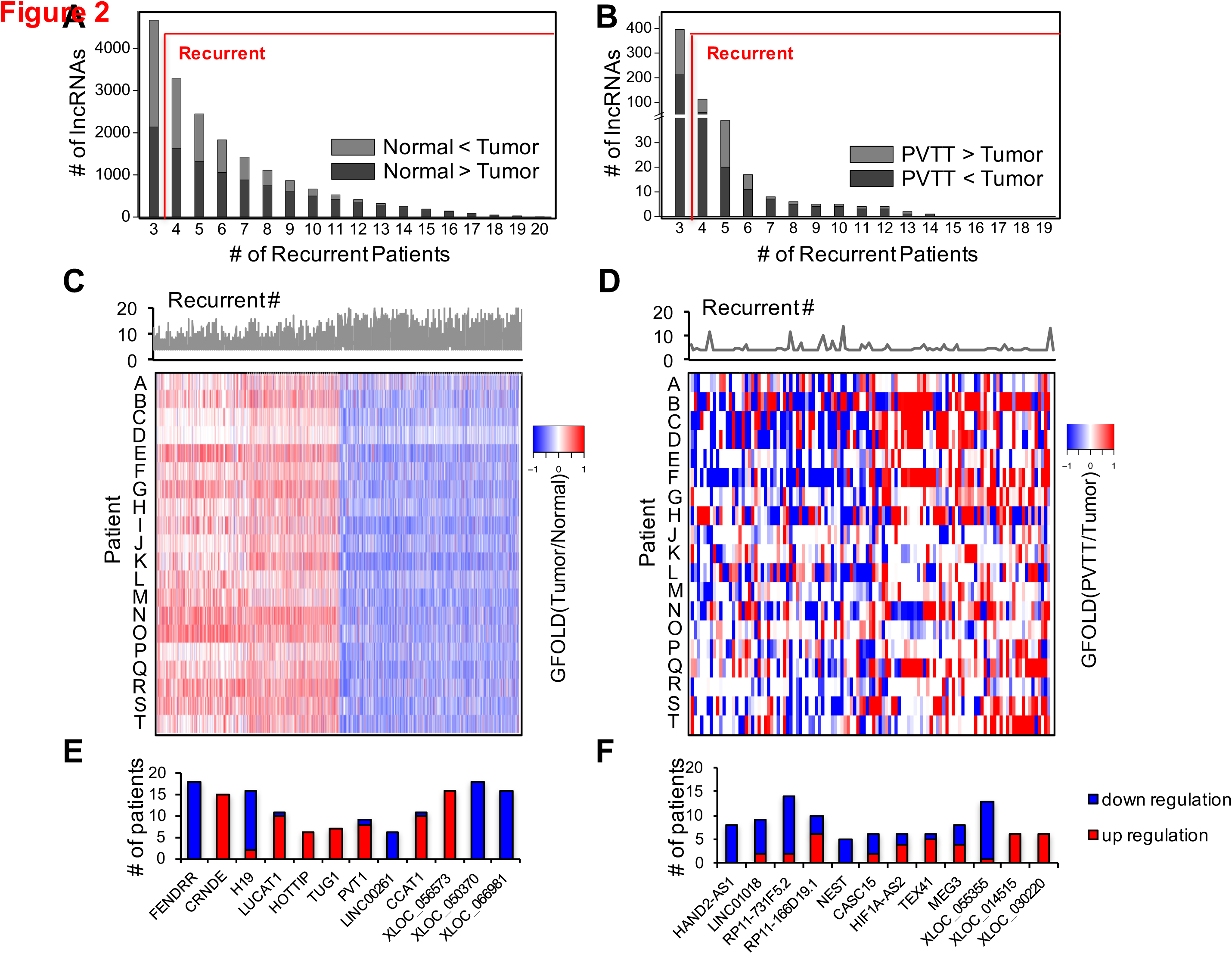
Recurrently deregulated lncRNAs in primary tumor tissue and PVTT. The recurrence of differentially expressed lncRNAs: (**A**) between adjacent normal tissue and primary tumor tissue, including normal > tumor and normal < tumor; (**B**) between primary tumor tissue and PVTT, including tumor > PVTT and tumor < PVTT. The x-axis is the number of patients in which the lncRNAs were differentially expressed. Recurrently deregulated lncRNAs were defined as those differentially expressed in at least 4 of 20 patients (20%). Fold-change of expression in each individual patient for (**C**) recurrently deregulated tumorigenesis-associated lncRNAs; (**D**) recurrently deregulated metastasis-associated lncRNAs (tumor V.S. PVTT). Patient I was not included in the analyses related to metastasis because the PVTT sample of patient I was contaminated. Stacked bar charts showing examples of recurrently deregulated lncRNAs, including tumorigenesis-associated (**E**) and metastasis-associated (**F**) lncRNAs. The number on the y-axis is the number of patients with differential expression of each lncRNA.

The fold-changes in expression (generalized fold change calculated by GFOLD (Feng et al. 2012)) of the recurrently deregulated lncRNAs (Figure 2C-D) and protein-coding genes were calculated (Supplementary Figure 7-8). The expression levels of each lncRNA and protein-coding gene in the PVTT samples showed much higher variance than those in the primary tumor samples. In addition, the numbers of recurrently deregulated lncRNAs (Supplementary Figure 6A-B) and protein coding genes (Supplementary Figure 8A-B) in PVTT samples from individual patients varied more than those in their primary tumor samples. Overall, expression levels of protein-coding genes and lncRNAs in PVTTs were much more heterogeneous than their expression levels in primary tumors, suggesting that more subtypes of individuals could exist with regard to metastasis than exist with regard to tumorigenesis.

### Examples of recurrently deregulated lncRNAs potentially associated with tumorigenesis and metastasis

Several of the tumorigenesis-associated lncRNAs (Figure 2E), including PVT1 and HOTTIP, were reported by previous studies. PVT1 promotes cell proliferation, cell cycle progression, and the development of a stem cell-like phenotype of HCC cells *in vitro*, while promoting HCC growth *in vivo* (Wang et al. 2014). A recent study reported that HOTTIP is significantly up-regulated in HCC specimens, associated with HCC progression, and predictive of clinical outcome (Quagliata *et al*. 2014).

Metastasis-associated lncRNAs (Figure 2F) were recurrently deregulated in the PVTT-tumor comparison. For example, several lncRNAs (TEX41, XLOC_014515, and XLOC_030220) were recurrently upregulated in PVTT samples, whereas others (HAND2-AS1, RP11-731F5.2, and XLOC_055355) were recurrently downregulated in PVTT samples. Some of the recurrently deregulated lncRNAs were also reported as deregulated in other cancer types. For example, HIF1A-AS2, a lncRNA antisense to hypoxia-inducible factor 1-alpha, is overexpressed in gastric cancer cells and involved in gastric cancer development (Chen et al. 2015b). Notably, we identified some newly assembled lncRNA candidates potentially related to metastasis, such as XLOC_014515 and XLOC_030220 (Supplementary File 6).

### Association of recurrently deregulated lncRNAs with clinical data in the TCGA and other published data

Based on Gene Set Enrichment Analysis (GSEA) (see Methods) of the recurrently deregulated lncRNA sets, we found that the expression levels of the tumorigenesis-and metastasis-associated lncRNA sets were consistent with their expression levels in another published dataset from 11 matched normal tissue samples, primary tumors, and PVTTs (Hong et al. 2015) (Figure 3A-B). In addition, we explored the expression levels of recurrently deregulated lncRNAs in a TCGA LIHC cohort (see Methods). Tumorigenesis-associated lncRNAs were also aberrantly expressed between normal tissues and primary tumors. Because the TCGA LIHC cohort had no PVTT or metastatic tumor samples, we compared expression levels of metastasis-associated lncRNAs between primary tumors with invasion and without invasion. We found that the deregulation of metastasis-associated lncRNAs was in accord with the clinical status of the TCGA patients (Figure 3A-B).

**Figure 3.**
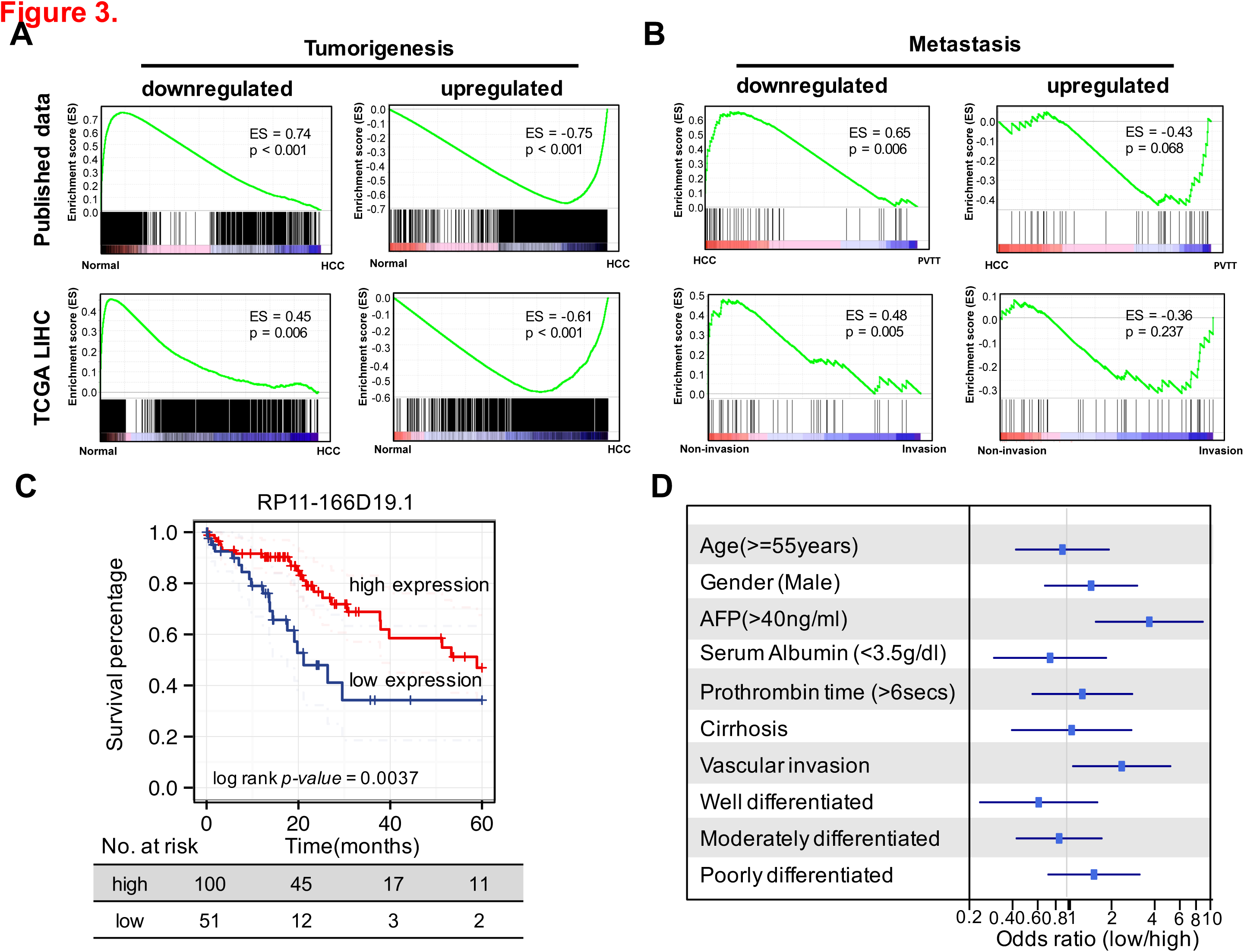
Association of recurrently deregulated lncRNAs with TCGA clinical data and other published data. (**A**) Gene set enrichment analysis (GSEA) of recurrently deregulated tumorigenesis-associated lncRNAs based on TCGA LIHC data and published liver cancer data. lncRNAs were rank-ordered by differential expression between adjacent normal tissue and primary tumor samples. (**B**) GSEA of recurrently deregulated metastasis-associated lncRNAs. lncRNAs were rank-ordered by differential expression between primary tumors with and without vascular invasion in TCGA LIHC data, as well as by differential expression between primary tumor tissue and PVTT in published liver cancer data. (**C**) Kaplan-Meier analysis of overall survival in the TCGA LIHC cohort. Subjects were stratified according to the expression of lncRNA RP11-166D19.1. The p-value for Kaplan-Meier analysis was determined using a log-rank test. (**D**) Multivariate analysis using other clinical information. Forest plot depicting the correlation between the indicated clinical criteria and the expression level of RP11-166D19.1.

The consistent expression levels of recurrently deregulated lncRNAs in our samples and the TCGA LIHC cohort suggest that the identified lncRNAs could potentially serve as biomarkers. As an example, we explored a metastasis-associated lncRNA, RP11-166D19.1 (ENSEMBL ID: ENSG00000255248, an isoform of MIR100HG, Supplementary Figure 9), which was also significantly downregulated in 4 of 11 PVTT samples in comparison with the paired primary tumors in the published liver cancer data (Hong et al. 2015). We found that RP11-166D19.1 could be used to clearly classify the patients in the TCGA cohort into two subclasses with different survival rates. RP11-166D19.1 expression was significantly associated with the overall survival time of HCC patients’ (Mantel-Cox test, log rank p-value = 0.0037) (Figure 3C). Moreover, a multivariate analysis based on additional clinical information showed that the HCCs of the low RP11-166D19.1 expression subclass were globally more severe than those of the high expression subclass: the low expression subclass was significantly correlated with AFP ≥ 40 ng/mL, had a high risk of vascular invasion, and had a clear tendency to be associated with serum albumin <3.5 g/dL and prothrombin time >6 secs; the patients in high expression subclass were mostly well differentiated (Figure 3D).

### DNA methylation levels and copy numbers of the recurrently deregulated lncRNAs are altered in HCC cells

We categorized the recurrently deregulated lncRNAs based on their correlation with copy number variations (CNVs) and/or DNA methylation alterations assayed in our matched samples (Figure 4A). We listed all recurrently deregulated lncRNAs (Supplementary File 7) with expression patterns correlated with CNV data (i.e., upregulated lncRNAs were found to be located in a DNA amplification region) and/or DNA methylation data (i.e., downregulated lncRNAs had high DNA methylation signals at their promoter regions). Several lncRNAs were recurrently (at least 4 patients) upregulated in some patients and recurrently downregulated in other patients; such lncRNAs were designated as bimorphic lncRNAs (Figure 4A).

The CNVs of recurrently deregulated lncRNAs in HCC cells were analyzed using CytoscanHD arrays. Based on a GISTIC analysis (Mermel et al. 2011), 4 significantly amplified genome regions and 70 significantly deleted genome regions were revealed in our samples (Figure 4B). To characterize candidate CNV-driven lncRNAs, we mapped recurrently deregulated lncRNAs to amplified and deleted genome regions. In total, 399 recurrently deregulated lncRNAs were identified in deleted regions, whereas none were found in amplified regions (Figure 4A, Supplementary File 7). For example, FENDRR, which was reported to be a prognostic biomarker in gastric cancer (Xu et al. 2014), had a similar pattern of decreased expression driven by copy number deletion (Figure 4B).

**Figure 4.**
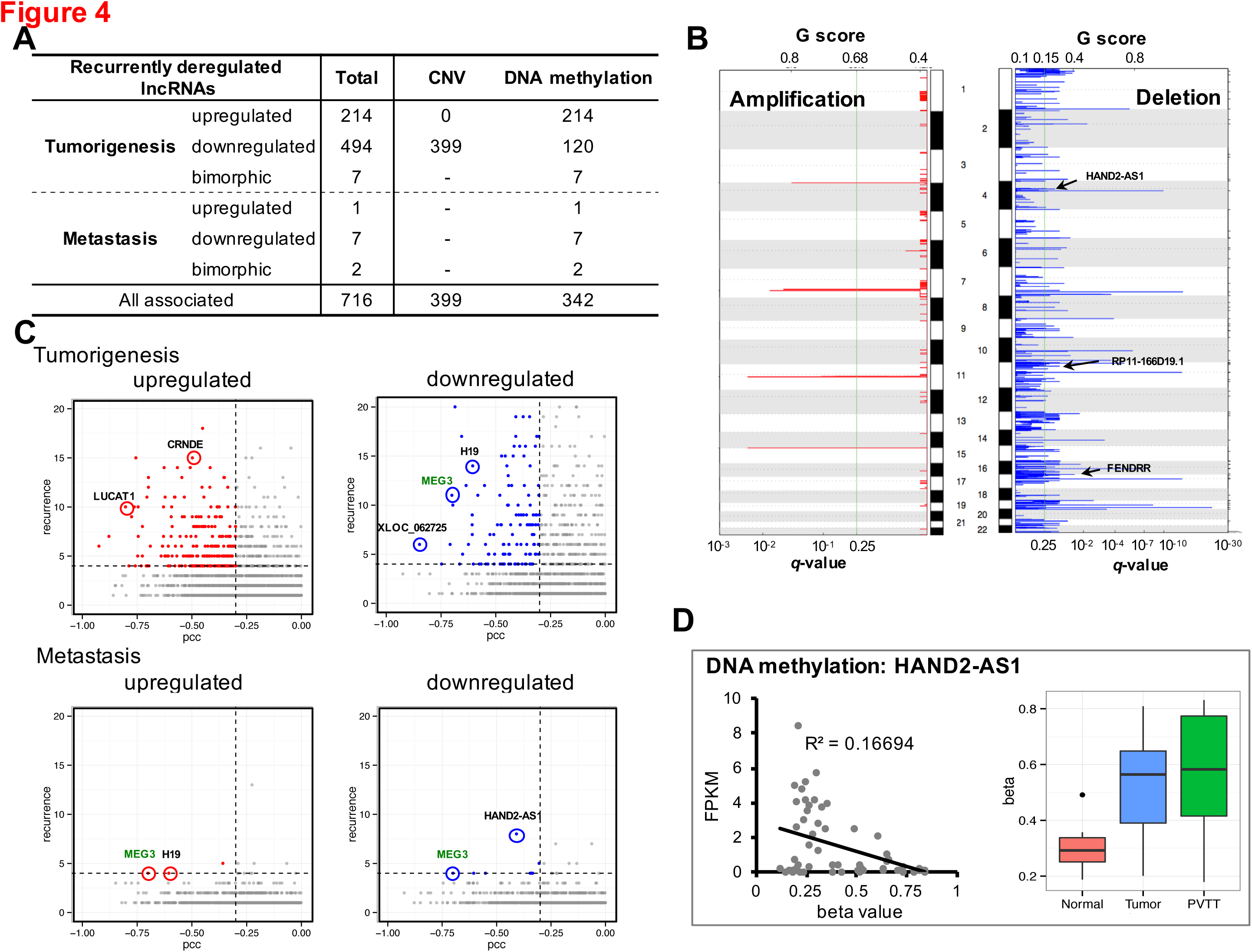
Regulation mechanisms for recurrently deregulated lncRNAs. (**A**) Summary of the regulation mechanisms of recurrently deregulated lncRNAs. The numbers of lncRNAs associated with CNV and/or DNA methylation data are listed. Bimorphic lncRNAs were those recurrently (>= 4 patients) upregulated in some patients and recurrently downregulated in other patients. The total is not equal to the sum because the overlaps between sub-types. (**B**) Chromosomal view of amplification and deletion peaks between primary tumor and normal tissue. The G-scores (top) and FDR q-values (bottom) of peaks were calculated using GISTIC2.0. The G-score considered the amplitude of the aberration and its frequency of occurrence across all samples. The q-value was calculated for the observed gain/loss at each locus using randomly permuted events as a control. Examples of recurrently deregulated lncRNAs located in the peaks (only found in deletions) are labeled. (**C**) Scatterplots showing recurrently deregulated lncRNAs that were putatively affected by alterations in DNA methylation. The upper and lower panels depict recurrently deregulated tumorigenesis-and metastasis-associated lncRNAs, respectively. Recurrently deregulated lncRNAs (recurrence >=4, y-axis) driven by DNA methylation had expression levels that were inversely correlated with DNA methylation levels at their promoter regions (PCC, Pearson correlation coefficients <-0.3, x-axis). Examples of recurrently deregulated lncRNAs are labeled. (**D**) Example of a recurrently deregulated lncRNA driven by DNA methylation. The HAND2-AS1 expression level (FPKM) was inversely correlated with its promoter methylation level (beta value). Boxplot showing that beta values in primary tumor samples were significantly higher than those in normal tissue samples, but slightly lower than those in PVTT samples.

Based on DNA methylation microarrays of 60 matched samples, we analyzed DNA methylation patterns to identify recurrently deregulated lncRNAs that were affected by alterations in DNA methylation. We applied several separate filtering criteria (see Methods) to define recurrently deregulated lncRNAs driven by alterations in DNA methylation. In total, 342 (10.7%) recurrently deregulated lncRNAs had significant correlations between DNA methylation and expression levels (Figures 4A and 4C, Supplementary File 7), suggesting that their expression levels in tumor and/or PVTT samples were probably regulated by DNA methylation. As an example, we showed that the expression level of a recurrently deregulated lncRNA, HAND2-AS1, was well inversely correlated with the DNA methylation level at its promoter region (R^2^ = 0.16694) (Figure 4D). The promoter region of HAND2-AS1 was hypermethylated in primary tumors and PVTT samples.

### Co-expression network infers potential metastasis pathways of the recurrently deregulated lncRNAs

To predict potential functional and regulatory mechanisms of lncRNAs with respect to the molecular etiology of HCC, we constructed a co-expression network (Liao et al. 2011) of protein-coding genes and lncRNAs (see Methods). The resulting co-expression network consisted of 7,367 protein-coding genes, 6,377 GENCODE lncRNAs, and 5,612 newly assembled lncRNAs. There were 1,286 recurrently deregulated lncRNAs in the network. All protein-coding genes and lncRNAs were grouped into 43 clusters, each of which had at least 100 highly inter-connected genes (Supplementary File 8). In addition to interaction edges within a cluster, there were also 337,609 edges between nodes in different clusters, which could indicate their functional relatedness or regulatory relationships (Figure 5A, Supplementary Figure 10).

**Figure 5.**
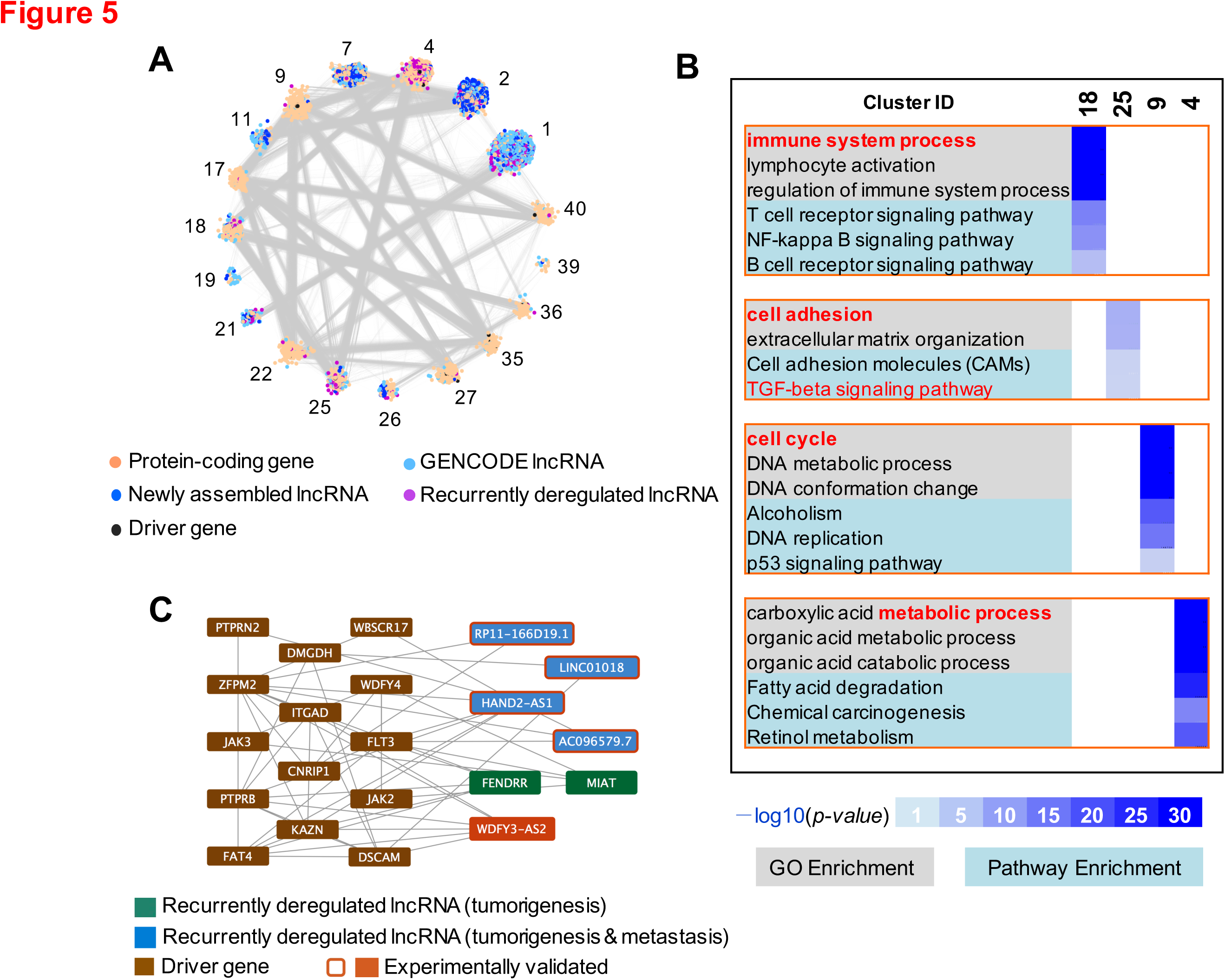
Co-expression network infers potential function pathways of recurrently deregulated lncRNAs. (**A**) Network representation of the 18 selected inter-connected clusters in the coding-non-coding co-expression network. (**B**) GO and KEGG pathway enrichment for 4 selected clusters (4, 9, 18, and 25). Heatmap showing the enrichment score (-log(*p-value*)) for various GO terms and KEGG pathways in 4 selected clusters. The most significantly enriched GO terms and KEGG pathways are displayed. (**C**) Sub-network showing important genes/lncRNAs in cluster 25. The subnetwork depicts the relationships among four lncRNAs and liver cancer-related driver genes. The network was drawn using Cytoscape (http://www.cytoscape.org/).

Among the 43 clusters, we found 4 clusters containing protein-coding genes with interesting functions: clusters 4, 9, 18, and 25 (Figure 5B). The recurrently deregulated lncRNAs are highly enriched in these gene clusters (Fisher’s exact test, all p-values <0.01). For example, Gene Ontology and KEGG pathway enrichment analyses suggested that the protein-coding genes in cluster 4 are mostly associated with metabolic processes in the liver, such as organic acid metabolism and degradation of fatty acids. The protein-coding genes in cluster 9 mainly function in cell cycle processes such as DNA repair, DNA replication, and DNA metabolism, which influence cell migration (Boehm and Nabel 2001). Cluster 18 is enriched with immune response genes involved in the T cell and B cell receptor signaling pathways, consistent with reports that immune and inflammatory responses play decisive roles in tumor development by influencing the processes of invasion and migration (Grivennikov et al. 2010).

Another intriguing example is cluster 25 (Figure 5C), which includes protein-coding genes enriched in functional terms related to metastasis, such as cell adhesion and the TGF-β signaling pathway, which have been shown to play essential roles in diverse processes, including cell proliferation, differentiation, motility, and adhesion (Padua and Massagué 2009). Furthermore, many HCC-related driver genes (Schulze et al. 2015) are also found in cluster 25. For example, the FLT3 receptor plays a role during the late stages of liver regeneration and is involved in GPCR signaling pathways (Aydin et al. 2007). FLT3 was co-expressed with other driver genes in the sub-network, including WDFY4 and FAT4. Moreover, four recurrently deregulated lncRNAs (HAND2-AS1, AC096579.7, FENDRR, and MIAT) were strongly correlated with FLT3 at the expression level, suggesting that these lncRNAs have functions related to that of FLT3. Another interesting gene in cluster 25, FAT4, encodes a cadherin (a calcium-dependent cell adhesion protein) that serves as a tumor-suppressor gene (Zang et al. 2012). FAT4 was closely associated with some recurrently deregulated lncRNAs, including HAND2-AS1, FENDRR, and WDFY3-AS2, all of which were differentially expressed during cell migration. Cell adhesion was one of the most significantly enriched processes during tumor metastasis. These co-expression relationships provide functional evidence demonstrating that adhesion-related lncRNAs likely have roles in tumor metastasis. Furthermore, driver gene ZFPM2 was also highly involved in the sub-network; it was significantly correlated with 7 driver genes and 5 recurrently deregulated lncRNAs. Some migration-related recurrently deregulated lncRNAs (HAND2-AS1, AC096579.7, and FENDRR) had expression patterns similar to that of PTPRB, which plays an important role in blood vessel remodeling and angiogenesis (Behjati et al. 2014), indicating that these lncRNAs could have related functions.

### Functional assay for candidate lncRNAs associated with metastasis

We have globally focused on the function of recurrently deregulated lncRNAs using a co-expression network. To confirm the putative candidate lncRNAs that might function in the progression of HCC, we used transwell migration assay to validate them. We first selected 10 lncRNA candidates that were associated with cell adhesion based on the co-expression network described above (Supplementary Table 2), in which 4 lncRNAs (WDFY3-AS2, HAND2-AS1, RP11-166D191, and XLOC_055355) were also significantly co-expressed with genes in the TGF-β signaling pathway. Note that 7 candidate lncRNAs (shown in Figure 6A) were recurrently deregulated lncRNAs in PVTT samples and designated metastasis-associated lncRNAs in the sections above. Three siRNAs were synthesized for each candidate lncRNA and mixed as a pool (Supplementary File 9). Three HCC cell lines, HepG2, SMMC-7721, and LM9, were used to conduct loss-of-function RNAi assays (Figure 6A).

**Figure 6.**
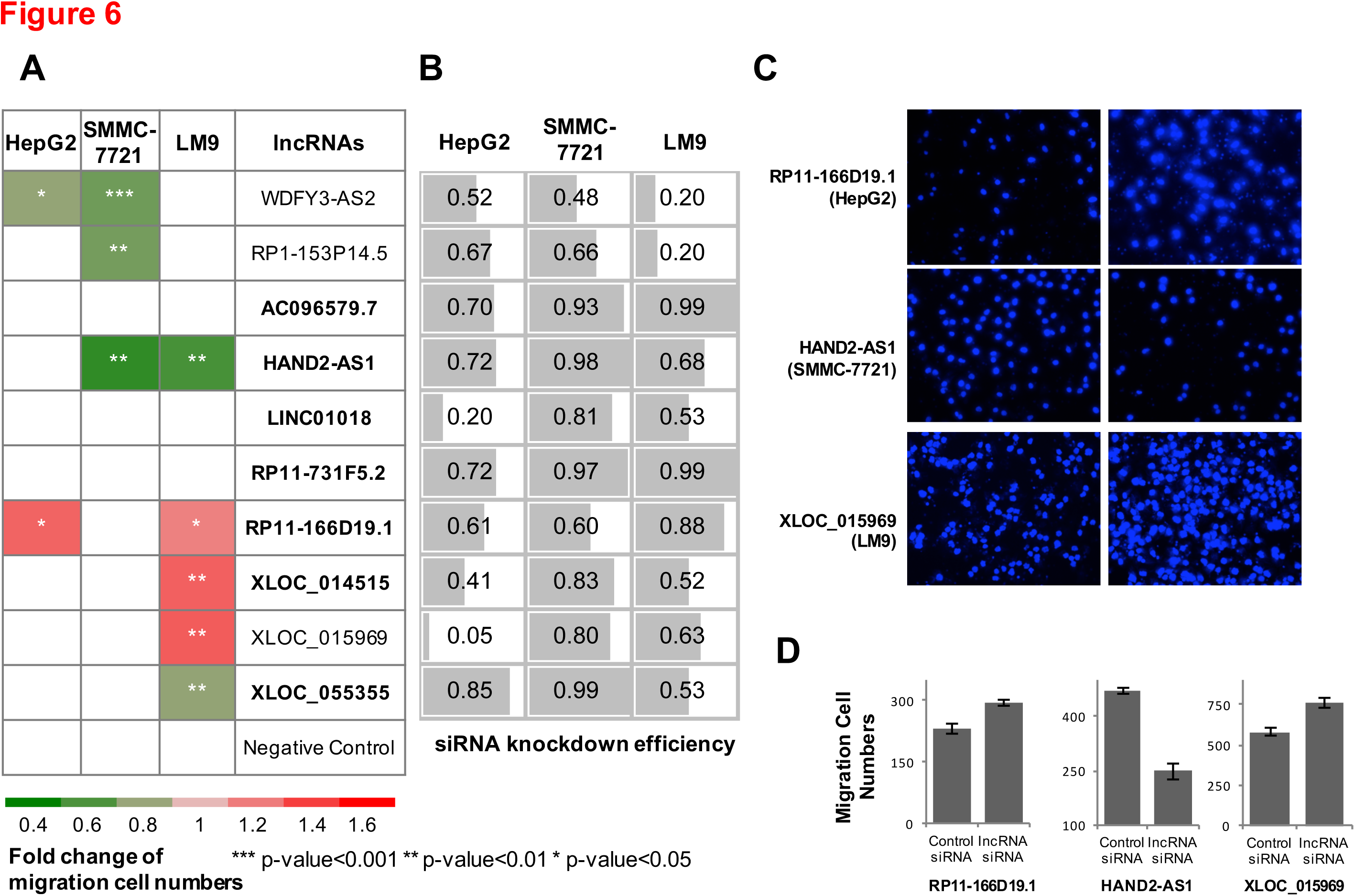
Loss-of-function assay of candidate lncRNAs regulating cell migration. Transwell migration assays were conducted to test the effects of siRNA-mediated RNAi of the candidate lncRNAs in three liver cancer cell lines: HepG2, SMMC-7721, and LM9. (**A**) The value in the heatmap is the fold-change (p-value <0.05) of the transwell cell numbers for knockdown cells over those of control cells. All results are expressed as the mean derived from three independent experiments. Student’s unpaired t-test (two-tailed) was used for comparisons of two groups. Seven of the ten candidate lncRNAs were metastasis-associated lncRNAs that were recurrently deregulated in PVTT (in bold font). (**B**) RNAi was validated by qRT-PCR. Examples of migration phenotype: transwell cells (DAPI staining) (**C**) and their counts (**D**).

Remarkably, knockdown of 7 of the 10 candidate lncRNAs significantly affected cell migration in at least one cell line; knockdown of 3 lncRNAs produced accordant alterations in cell migration in at least two cell lines by suppressing or promoting cell migration (Figure 6A, Supplementary File 9). RNAi knockdown efficiency was confirmed using qRT-PCR (Figure 6B, Supplementary Figure 11). The results from three representative transwell assay are shown on the right side of the figure (Figure 6C,D). For some lncRNAs knockdown experiments, changes in invasion ability were in consistent with the HCC patients’ deregulation patterns. For example, RP11-166D19.1 was recurrently downregulated in PVTT samples from 4 patients. The loss-of-function assay showed that knockdown of RP11-166D19.1 further increased the migration ability of HCC cells. However, some other lncRNAs, such as HAND2-AS1, demonstrated an inconsistent trend between deregulation patterns in HCC patients and experimentally validated functions in cancer cell lines; silencing of HAND2-AS1 suppresses cell migration, although it was downregulated in 8 of 20 patients’ PVTT samples. It has been reported that nearly half of HCC cell lines do not resemble primary tumors (Chen et al. 2015a), so the intrinsic differences between cancer cells lines and clinical samples might explain the discrepancy between the samples’ gene expression patterns and experimentally validated functions in cell lines. Overall, the high validation rate of the candidate lncRNAs showed that the co-expression network, based on previous knowledge of signaling pathways and supplemented by recurrent aberrant expression patterns in matched clinical samples, identified functional lncRNAs involved in the sophisticated regulation of cancer development and progression.

## Discussion

Based on genomic, epigenomic, and transcriptomic data of HCC primary tumors and PVTTs, this study reports several innovative findings. First, based on high-throughput sequencing technology and bioinformatics analysis of 60 matched samples, including primary tumors, PVTTs, and adjacent normal tissue, we discovered and characterized an expanded landscape of known and novel lncRNAs relevant to HCC. Moreover, we identified lncRNAs that were recurrently deregulated during HCC tumorigenesis and metastasis. Second, integrative multi-omics analysis revealed that recurrent deregulation of lncRNA expression was often associated with alterations in DNA methylation and CNV. In addition, lncRNA expression levels were correlated with clinical data from the TCGA and other published liver cancer data. Lastly, using network analysis and loss-of-function assays, we identified functional lncRNAs potentially related to cell adhesion, immune responses, and metabolic processes. For example, our paired RNA-seq data showed that lncRNA HAND2-AS1 was recurrently deregulated; its expression levels among 60 samples were inversely correlated with the matched DNA methylation data. Then, based on our co-expression network, we inferred that HAND2-AS1 might be related to metastasis. Finally, using an RNAi functional assay, we demonstrated that the function of lncRNA HAND2-AS1 in HCC cells is related to cell migration.

In addition, we have shown that RP11-166D19.1 could potentially serve as a promising single-gene biomarker of HCC. We also demonstrated that knockdown of RP11-166D19.1 promoted cell migration. RP11-166D19.1 is an isoform of lncRNA MIR100HG (Supplementary Figure 9), a leukemia-related oncogene (Emmrich et al. 2014) hosting three miRNAs (let-7a, miR-100, miR-125b) in its introns (Chang et al. 2015). As reported previously, lncRNAs are more tissue-and cell-type-specific than protein-coding genes (Cabili et al. 2011). The local secondary structures of lncRNAs make them more stable and provide a greater likelihood of detection (Yan et al. 2015). Therefore, translation of these results into novel lncRNA biomarkers might impact clinical decision-making and ultimately improve clinical outcomes for patients with HCC.

By exploring lncRNA transcriptome alteration, we found that the lncRNA landscapes of PVTTs were indistinguishable from those of matched primary tumors, consistent with previous studies (Ye et al. 2003). We employed principal component analysis (PCA) to assess the expression profile of recurrently deregulated lncRNAs in different samples. PCA results showed that the recurrently deregulated lncRNAs could clearly separate the primary tumors from adjacent normal tissues, while the PVTTs were more similar to the primary tumors (Supplementary Figure 7). This observation showed that the lncRNA expression profile of the PVTTs was very similar to that of their matched primary tumors, consistent with studies on protein-coding genes, CNV, and DNA methylation (Ye et al. 2003; Gu et al. in submission). The above data suggest that 1) primary tumors of HCC patients with PVTT may contain sub-clones with the potential to invade the portal vein and develop into PVTTs; 2) many metastasis-associated lncRNAs were deregulated in these sub-clones. These findings are consistent with clinical observations, because all of our sequenced patients had stage IV HCC with serious PVTT. On the other hand, although the overall lncRNA expression patterns of PVTTs were similar to those of their matched primary tumors, approximately 100 lncRNAs were significantly and recurrently deregulated in PVTTs, compared with the paired primary tumors. The results suggest that these lncRNAs could play essential roles in metastasis, because they were deregulated further as primary HCC cells invaded the portal vein.

Overall, we identified tumorigenesis-and metastasis-associated lncRNAs recurrently deregulated, many of which have been validated by the followed assays and mechanistically linked to cancer development and progression. We anticipate that the recurrently deregulated lncRNAs identified in this report could provide a valuable resource for studies aimed at delineating the relationship between functional lncRNAs and HCC tumorigenesis/metastasis.

## Materials and Methods

### RNA-seq data and transcriptome assembly for 60 samples from HCC patients

Total RNA from 60 samples from 20 Chinese HCC patients was sequenced (GSE77509). Each patient had three matched samples: primary HCC tumor, adjacent normal liver tissue, and portal vein tumor thrombosis (PVTT). The patients were ordered using alphabetic labels (A to T) in this paper, but the patients were originally numbered as 3,6, 7, 8, 10, 11, 12, 13, 14, 15, 16, 17, 18, 19, 20, 21, 22, 24, 25, and 26 (Gu et al. in submission). The PVTT sample of one patient (14) was not distinguishable from normal tissue(Gu et al. in submission)), so we did not use it in our migration and metastasis analyses.

We first evaluated RNA-seq quality using FastQC (version 0.10.1) and found that all raw reads qualified for the analysis. We aligned the RNA-seq reads to human reference rRNA using Bowtie with one mismatch in order to estimate the rRNA ratio. Most of the rRNAs were removed by our experiments; only a few remained and generated rRNA reads.

Next, the RNA-seq reads were mapped to the human reference genome (hg19) using Tophat (version 2.0.10) (Trapnell et al. 2009) with default parameters. The human genome sequence was downloaded from Ensembl (*Homo sapiens* GRCh37/hg19). After mapping, we further removed the PCR duplicates using *rmdup* in Samtools (Li et al. 2009). Further details of the preprocessing results are described in Supplementary File 1.

Subsequently, based on the mapped reads, we re-assembled a transcriptome using Cufflinks (version 2.2.1) (Trapnell et al. 2010) by providing reference annotations (option “-g”) from GENCODE (v19) for each dataset of 60 samples. Next, we used Cuffmerge (Trapnell et al. 2010) to merge all 60 meta-assemblies to generate a final transcriptome (Supplementary File 2).

### Identification of the newly assembled lncRNAs

After the final transcriptome was constructed, we used several stringent filters to identify a set of lncRNAs:

[1] Transcripts that overlapped (>=1 nt) on the same strand with the exons of protein-coding genes or noncoding RNAs (both canonical ncRNAs and long lncRNAs (lncRNAs)) annotated by GENCOE (V19) were removed. Canonical ncRNAs include rRNA, tRNA, miRNA, snRNA, snoRNA, misc_RNA, mitochondria tRNA and rRNA. Six biotypes were defined as “long non-coding RNAs” by GENCODE: “lincRNA”, “processed_transcript”, “sense_intronic”, “sense_overlapping”, “antisense”, and “3prime_overlapping_ncrna” (Harrow et al. 2012). Note that “processed transcript” means that they do not contain an open reading frame (ORF), although they could have protein-coding-style names historically. Note that some lncRNAs were updated as protein-coding genes in recently released GENCODE annotation versions.
[2] Transcripts shorter than 200 bp and without strand information were discarded.
[3] Single-exon transcripts proximal (within 2000 bp) to protein-coding genes or other noncoding RNAs on the same strand were filtered.
[4] Expression levels for single-exon transcripts were calculated using Cufflinks (Trapnell et al. 2010) with rRNA reads masked (FPKM must have been greater than 0.5 in at least one sample).
[5] A coding potential calculator, CPC (Kong et al. 2007), was used to predict the transcripts’ coding potential with default parameters. Transcripts with CPC score >0 were excluded. To ensure a stringent lncRNA definition, we used our noncoding RNA prediction model, *RNAfeature* (Hu et al. 2015), to further remove remaining transcripts with coding potential.

### Annotation of lncRNAs

We annotated the genomic locations of the identified lncRNAs, as well as GENCODE and MiTranscriptome lncRNAs (TCGA), by overlapping them with annotated coding genes. Intronic lncRNAs were defined as those located in the intronic regions of coding genes on the sense strand. Antisense lncRNAs overlapped with any exon (including both coding genes and ncRNAs) on the antisense strand with at least 1 nt. Cis-lncRNAs (also called sense lncRNAs) were those lncRNAs close to (within 2000 nt of the 5′ or 3′ ends) a protein-coding gene. The remaining lncRNAs that did not overlap with any coding genes or annotated ncRNAs were designated intergenic lncRNAs.

We also studied the overlap of the lncRNAs with pseudogenes and transposable elements because previous studies suggested that some lncRNAs could be derived from them. Annotations of pseudogenes and transposable elements were derived from GENCODE and the UCSC Genome Browser, respectively.

Furthermore, we also annotated lncRNAs with domains/motifs conserved with annotated canonical ncRNAs at the sequence and structure levels. Sequence conservation was based on performing BLASTN over the canonical ncRNAs’ sequences. The cutoff E-value was the default value of 1e-5. Secondary structure conservation was calculated by scanning the Rfam’s structure families of known ncRNAs using INFERNAL/cmscan (cutoff of the E-value was 0.01), in which hits are considered to be reliable enough to be reported in a possible subsequent search round.

### Conservation and SNP enrichment analysis for lncRNAs

We downloaded the PhastCons scores for multiple alignments of 46 vertebrate genomes from the UCSC Genome Browser (https://genome.ucsc.edu/). We then calculated two conservation scores for each transcript: one based on the average value of PhastCons scores in the exonic regions and the other based on those in the intronic regions.

To assess SNPs in different genomic elements, we downloaded two SNP databases: 1) 12,891 SNPs from the National Human Genome Research Institute’s GWAS catalog (https://www.genome.gov/26525384);2) 14,416,369 common SNPs from dbSNP Build 142 common (downloaded from the UCSC Genome Browser) (treated as background variation). We calculated the number of SNPs that overlapped with the transcripts using the BEDTools *intersect*function. We first calculated the fraction, *frac.(transcripts)*, of the amount of overlapped SNPs from the GWAS catalog to the number of overlapped background SNPs for different categories of genomic elements (e.g., lncRNAs and protein-coding genes). Next, we shuffled the transcripts’ positions on the whole genome 100 times and re-calculated *frac.(shuffled transcripts)*. Subsequently, we calculated the odds ratio (OR) as

OR = *frac.(transcripts)/frac.(shuffled transcripts)*.

An OR (control) was calculated by replacing SNPs from the GWAS catalog with control SNPs:

OR (control) = *frac.(transcripts)’/frac.(shuffled transcripts)’*, where the control SNPs were randomly selected from the background SNPs shuffled over the whole genome. The significance of enrichment for the OR over OR (control) was tested via paired Student’s t-test.

### Expression abundance and differential expression analysis for lncRNAs

We identified differentially expressed genes/lncRNAs between primary tumor samples and matched adjacent normal tissue, as well as between PVTT and matched primary tumor samples, for each individual patient. The differential expression analysis was conducted with GFOLD (version1.1.3) (Feng et al. 2012), because it was claimed to be a robust method of assessing samples without replicas. The significance cutoff of GFOLD was set at 0.01 (option:-sc 0.01). Genes/lncRNAs with fold-changes greater than two-fold were selected to independently define tumorigenesis-associated genes/lncRNAs and metastasis-associated genes/lncRNAs for each patient. In the supplementary files, expression abundance (FPKM calculated using Cufflinks) is provided (Trapnell et al. 2010).

### Kaplan-Meier survival, multivariate analysis, and GSEA based on TCGA LIHC samples’ clinical data

We downloaded RNA-seq data from 157 LIHC (liver hepatocellular carcinoma) patients with clinical data in the TCGA from the NCI’s Cancer Genomics Hub (CGHub) (Wilks et al. 2014). We calculated each gene/lncRNA’s expression level (FPKM) for each TCGA sample using Cufflinks (Trapnell et al. 2010).

In the Kaplan-Meier survival analysis, the survival data included vital status, days to death, etc., which were available for 151 of 157 patients. We first divided the samples into two groups (51 low expression and 100 high expression) according to a marker gene/lncRNA’s (e.g., RP11-166D19.1) expression level. Next, we used Kaplan-Meier survival analysis (Efron 1988) to perform a 5-year survival analysis via the *survival* package (https://cran.r-project.org/web/packages/survival) in the R environment for statistical computing and computed significance using the log-rank test.

Other clinical information (age, gender, AFP, serum albumin, prothrombin time, cirrhosis, vascular invasion, etc.) for the LIHC patients in the TCGA was downloaded for the multivariate analysis. Based on two groups defined by the expression level of a particular lncRNA (e.g., RP11-166D19.1), the odds ratio of each clinical criterion was calculated for each class of patients (low expression and high expression). A forest plot was drawn with odds ratios and 95% confidence intervals for each clinical criterion.

We used Gene Set Enrichment Analysis (GSEA v2.0.13) (Subramanian et al. 2005) to assess enrichment of sets of recurrently deregulated lncRNAs in other data sets. GSEA requires three input files: a gene set, expression data, and phenotype labels. We used the recurrently deregulated lncRNA set (the tumorigenesis set or metastasis set) as the gene set. Expression data were derived from the TCGA cohort or published liver cancer data (Hong et al. 2015). For the published data, we used the sample information (adjacent normal tissue, primary tumor, and PVTT) as phenotype labels. Because the TCGA LIHC cohort had no PVTT or other metastasis samples, we classified primary tumor samples into invasion and non-invasion groups based on clinical information (T stages in the TNM staging system: T1 versus T2-T4). The lncRNAs were rank-ordered by differential expression (signal2Noise in GSEA v2.0.13) (Subramanian et al. 2005) between the two groups.

### Copy number variation data for lncRNAs

DNA copy numbers (GSE77275) were probed for the 60 matched samples (PVTT/tumor/normal tissue samples from 20 patients) using Affymetrix CytoscanHD arrays following the manufacturer’s protocol (Gu et al. in submission). The CytoscanHD array contains 2,696,550 probes, including 1,953,246 nonpolymorphic probes. The GISTIC algorithm (GISTC2.0) (Mermel et al. 2011) was used to calculate G-scores and false discovery rates (q-values) for the aberrant regions and thus identify genomic regions that were significantly amplified or deleted across all samples. G-scores consider the amplitude of the aberration and its frequency of occurrence across all samples. Aberrant regions were considered significant if the assigned FDR q-value was less than 0.25. The GISTIC algorithm also reported genes found in each aberrant region. We identified recurrently deregulated lncRNAs for which copy number variations contributed to their deregulation.

### DNA methylation data for lncRNAs

DNA methylation profiles (GSE77269) were probed using the Illumina Infinium HumanMethylation 450k BeadChip (Gu et al. in submission), which contains more than 485,000
CpG sites. β values were calculated to independently assess the methylation levels of the CpG sites for each dataset. CpG sites were distributed across the promoter or gene body of the lncRNAs. To identify recurrently deregulated lncRNAs that were epigenetically regulated by DNA methylation, we assigned all CpG sites corresponding to promoter regions (2000 bp upstream of the transcription start site) to each lncRNA. Pearson correlation coefficients between expression levels and β values were calculated for each lncRNA and all assigned methylation sites across all 60 samples. When there were multiple CpG sites for the same gene promoter, the CpG site with the highest correlation was assigned to that lncRNA. Recurrently deregulated lncRNAs with Pearson correlation coefficients <-0.3 were identified as lncRNAs regulated by DNA methylation alteration.

### Co-expression network construction for lncRNAs and protein-coding genes

We adapted a published method (Liao et al. 2011) to construct a co-expression network of lncRNAs (including both GENCODE lncRNAs and lncRNAs identified in the HCC samples). The expression levels derived from the total RNA-seq data for the 60 samples were used. Genes whose maximum expression level among all the datasets ranked in the bottom 20% were excluded from the input gene list. For each gene pair (including lncRNAs and protein-coding genes), we calculated the Pearson correlation coefficient and the corresponding Student’s t-test p-value using the *WGCNA* package for the R environment for statistical computing (Langfelder and Horvath 2008). All p-values were adjusted for multiple testing via the Bonferroni correction in the *multtest* R package (Pollard et al. 2008). Markov clustering (MCL) (Enright et al. 2002) was used to detect highly inter-connected gene/lncRNA clusters. Bonferroni adjusted p-values (cutoff: 0.01) were used as edge weights for MCL. To control the size of the clusters generated from the MCL clustering, 2.4 was used as the inflation coefficient.

### Gene ontology and pathway enrichment analysis

For the protein-coding genes in each co-expression cluster, we used R package *topGO* (Alexa and Rahnenfuhrer 2010) to estimate enrichment in biological process (BP) terms for different gene sets. We estimated the significance of GO term enrichment using a hypergeometric test. Moreover, we used R package *KEGGREST* (Tenenbaum 2013) to estimate enrichment in biological pathways for each cluster. We also annotated 271 driver genes of liver cancer, which were derived from a recent study (Schulze et al. 2015).

### Function study of candidate lncRNAs using the transwell migration assay

Three siRNAs were designed for each candidate lncRNA (Supplementary File 9). qRT-PCR was used to monitor siRNA knockdown efficiency. Transwell migration assays were used to test the candidate lncRNAs’ effects on cell migration. Cells were transfected with lncRNA siRNAs (GenePharma) for 48 h, followed by resuspension and washing with phosphate-buffered saline (PBS) buffer. Cells (40,000) were seeded into the upper chamber of a transwell insert (pore size, 8 μm, Costar) in 100 μL of serum-free medium per well. Medium (600 μL) containing 10% serum was placed in the lower chamber to act as a chemoattractant. HepG2, LM9, and SMMC-7721 cell lines were each incubated for 24 h, after which nonmigratory cells were removed from the upper chamber by scraping with cotton. The cells remaining on the lower surface of the insert were fixed with 4% formaldehyde (Sigma) and stained with DAPI for counting. Each type of cell was assayed in triplicate.

## Data access

The high-throughput sequencing data from this study have been submitted to the NCBI Gene Expression Omnibus (GEO, http://www.ncbi.nlm.nih.gov/geo/) under accession number GSE77276 (RNA-seq, GSE77509; miRNA-seq, GSE76903; 450k array, GSE77269; Cytoscan HD array, GSE77275).

## Acknowledgements

This work was supported by National Key Basic Research Program of China [2012CB316503]; National High-Tech Research and Development Program of China [2014AA021103]; National Natural Science Foundation of China [31271402, 31522030]; Tsinghua University Initiative Scientific Research Program [2014z21045]. This work was also supported by Computing Platform of National Protein Facilities (Tsinghua University). The funding for open access was provided by National Key Basic Research Program of China [2012CB316503]. The results published here are in whole or part based upon data generated by The Cancer Genome Atlas managed by the NCI and NHGRI. Information about TCGA can be found at http://cancergenome.nih.gov.

## URLs

GENCODE, http://www.gencodegenes.org/

ENSEMBL, http://www.ensembl.org/

NHGRI GWAS catalog, https://www.genome.gov/gwastudies/

CGA, http://cancergenome.nih.gov/

ICGC, https://icgc.org/

CGHub, https://cghub.ucsc.edu/

